# Proteogenomic characterization reveals antitumor mechanisms of intrahepatic cholangiocarcinoma

**DOI:** 10.1101/2023.08.14.553174

**Authors:** Weibing Wang, Shifeng Xu

## Abstract

Diagnosing and managing intrahepatic cholangiocarcinoma (iCCA) poses a significant oncological hurdle as it is a deadly form of liver cancer.A significant role is played by nonapoptotic regulatory cell death (NRCD) in the tumor immune microenvironment (TME) of iCCA. Nevertheless, the precise functions of NRCD-associated genes (NRGs) in the tumor microenvironment (TME) are still not well understood. Transcriptomics, proteomic analysis, and single-cell RNA analysis were utilized to distinguish two NRG-related clusters in iCCA patients from the FU-iCCA cohort in this study. We have shown the clear disparities in immune traits and predictive categorization among two groups. To address the risk stratification and prognosis prediction, a prognostic signature model called NPS (NRG-related risks core prognostic signature model) was created using the FU-iCCA cohort. The validation of the NRG-associated risk score in prognosis and immunotherapy confirmed its predictive capabilities. In iCCA patients with high-risk, the secretion of CP and TGF-β proteins strengthened the enriched TGF-β signaling network between CD4+ T cells and erythroid cells. Patients exhibiting a low-risk score demonstrated improvement in the effector function of CD4+ T cells, resulting in a more favorable reaction to chemotherapy medications. The NPS-risk score showed a significant negative correlation with the IC50 values of four drugs (Trametinib, CI-1040, X17-ACC, and PD-0325901). From these findings, it can be inferred that a clear connection exists between the NRG and the TME in iCCA. Additionally, the risk score has the potential to act as a reliable prognostic indicator, offering advantageous outcomes for chemotherapy and immunotherapy. This could aid in making informed clinical decisions for patients with iCCA.

## Introduction

Intrahepatic cholangiocarcinoma (iCCA) is a form of primary liver cancer that has a high occurrence rate, ranking second only to hepatocellular carcinoma (HCC), which develops from the epithelial cells of the intrahepatic bile duct^1^. iCCA is not as prevalent as HCC, but its incidence rates have been increasing rapidly^2^. It is an extremely aggressive cancer, and long-term survival is only possible in patients who undergo a complete R0 surgical resection. Possible factors that increase the risk include long-term hepatitis and cirrhosis, inflammatory diseases affecting the bile ducts, and parasitic infections in the liver and bile ducts. It is important to note that in the majority of instances, the specific risk factor remains unidentified. The occurrence of iCCA in patients with primary sclerosing cholangitis is estimated to be between 5% and 10%, making it a known risk factor for iCCA development^3^. The pathophysiology of iCCA is intricate, with numerous signaling pathways implicated. For instance, long-term damage and swelling of the bile ducts lead to the activation of nitric oxide synthase as a result of cytokine generation.The enzyme will generate nitric oxide (NO), which will stimulate the expression of cyclooxygenase-2. This activation of cyclooxygenase-2 will then trigger the activation of various growth factors, resulting in cellular growth^4^. Furthermore, NO has the ability to directly oxidize and harm DNA, resulting in the occurrence of mutagenesis^5^. Pathogenesis could also be influenced by epigenetic alterations. Unlike other mutations, the mutations found in isocitrate dehydrogenase (IDH)1 and 2 are detected in approximately 18% of iCCAs, making them particularly associated with intrahepatic forms of cholangiocarcinoma^6^. There have been reports of specific mutations being linked to hepatitis B surface antigen (HBsAg)^7^. In the past, it was often misdiagnosed as HCC. Diagnosing, staging, and managing iCCA has become increasingly challenging due to the significant oncological hurdle of a 38% five-year survival rate for patients with iCCA.

Non-apoptotic regulatory cell death (NRCD) can be subdivided into autophagy, ferroptosis, pyroptosis, necroptosis, and cuproptosis ^8,9^. And research on apoptosis has been conducted for more than 30 years. However, therapeutic agents targeting apoptosis regulators such as apoptosis-associated cystathionases or B-cell lymphoma 2 (BCL-2) family proteins have been ineffective in antitumor therapy^10^. In contrast, NRCD is significantly associated with cancer progression and response to therapy. For example, photodynamic therapy (PDT), a potential therapy for iCCA, has the ability to regulate ferroptosis by generating reactive oxygen species (ROS)^11^. Ferroptosis can also enhance the metabolic and inflammatory regulation of tumor-associated macrophages to stimulate effective tumor-killing activity^12^. There might be a promising approach for the treatment of iCCA, targeting SHARPIN-mediated cell ferroptosis via the p53/SLC7A11/GPX4 signaling pathway. Furthermore, inhibition of autophagy by targeting PIKfyve enhances the response to immune checkpoint blockade in prostate cancer^14^; and inhibition of Aurora Kinase A induces necroptosis in pancreatic cancer to slow in situ tumor growth in mice^15^. Song et al. determined the therapeutic liability of pyroptosis-related genes in targeted therapy and immunotherapy for colorectal cancer ^16^. These have been quite surprisingly effective. Thus, targeting NRCDs may be a new light in the treatment of cancers, including iCCA.

Recent progress in genomic and transcriptomic sequencing analysis has revealed the genetic landscape of iCCA. The multi-dimensional ‘‘omics’’ strategies encompassing proteomic and phosphoproteomic profiling in conjunction with genomic analysis, which may help reveal new mechanisms and identify novel targets for developing therapies to provide additional treatment options for iCCA patients ^17^. For the purpose of revealing the new mechanisms, we have analyed the information of 207 Chinese iCCA patients from the FU-iCCA cohort to performe an integrative genomic, transcriptomic, proteomic, and phosphoproteomic.

In our study, NRCD-related clusters were identified based on the FU-iCCA cohort by utilizing the unsupervised clustering algorithm. Based on the 10 seven NRCD-related genes (NRGs), which is screened out using the LASSO Cox regression model, an NRG prognostic signatures (NPS) model for iCCA patients was constructed from the FU-iCCA cohort. We have compared the difference between two NRG-related clusters and between high- and low-risk groups in enriched annotation, TME and immunotherapy response. Its prediction abilities of survival probability and treatment effect of immune checkpoint inhibitor, and chemotherapy were evaluated in different platforms datasets. Finally, a nomogram was constructed to quantify survival probability combined with the risk score and other prognostic clinical features. Our integrated proteogenomic analysis illustrated that genetic alterations may bring about phenotypic perturbations. Meanwhile, it also revealed disease subgroups with distinct features and prognosis, which delineated the complex mechanisms of iCCA pathogenesis for prospective exploration of precision management.

## Methods

### Data sources

Figure 1 have showed a map of the process of the present work. Gene expression (fragments per kilobase million, FPKM) and the relevant prognostic and clinicopathological data of iCCA have been downloaded from the FU-iCCA cohort ^17^ and The Cancer Genome Atlas (TCGA) (https://portal.gdc.cancer.gov/) database. We have downloaded the information about IMvigor210 cohort including complete expression data and detailed clinical information of patients with metastatic urothelial cancer treated with anti-PD-L1 agents from the R package IMvigor210 CoreBiologies (ver-sion1.0.0) ^18^. GSE78220 contains survival information of iCCA patients treated with anti-PD/anti-PD-L1.

**Figure 1.**
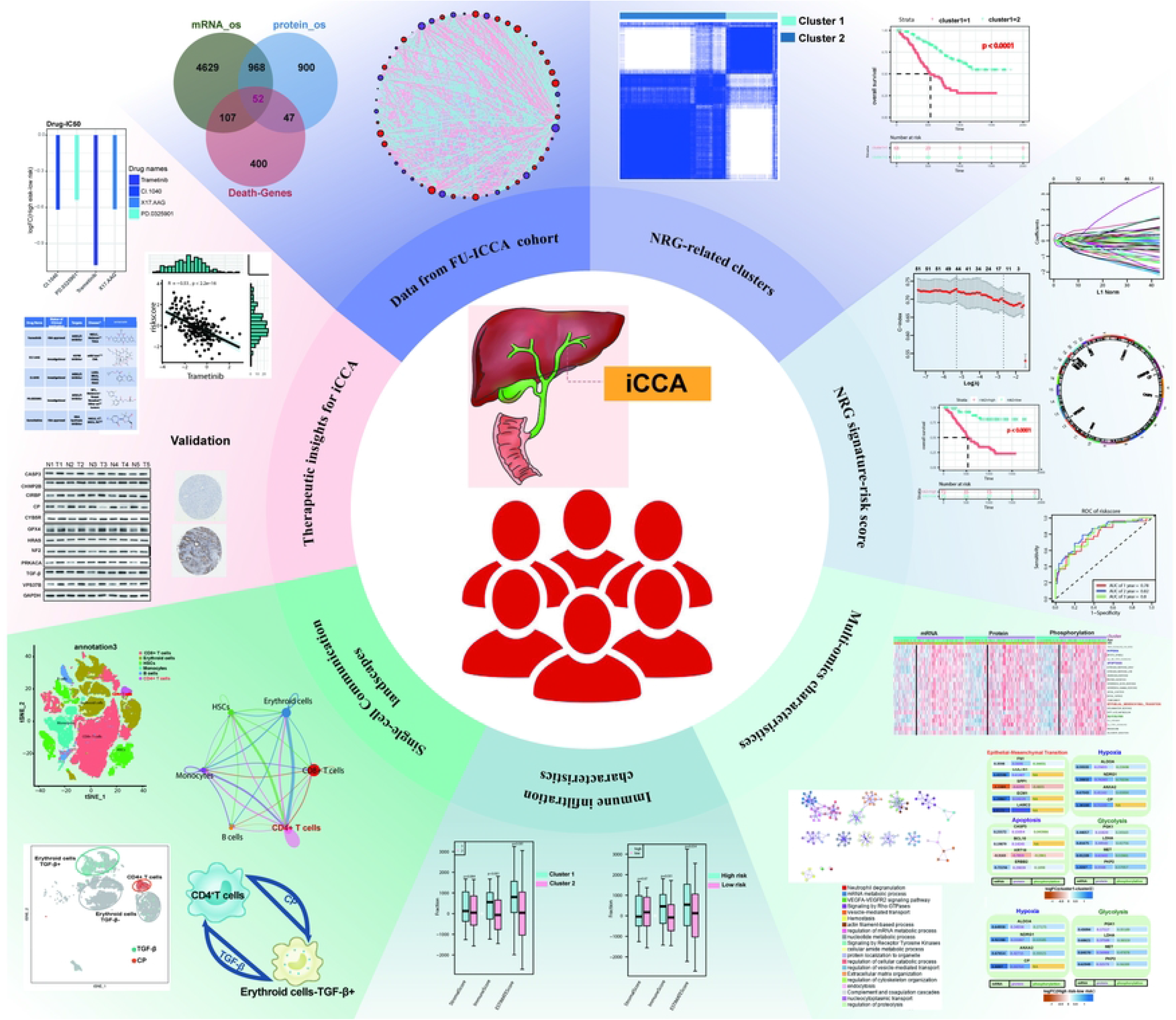
The entire analytical process of the study.

The nonapoptotic regulatory cell death (RCD) contains necroptosis, pyroptosis, autophagy, cuproptosis and ferroptosis. In our study, we have obtained several information, which contains necroptosis, pyroptosis, autophagy-related genes, from MsigDB Team (Necroptosis-KEGG, Pyroptosis-REACTOME, Autophagy-KEGG) (http://www.broad.mit.edu/ gsea/msigdb/). Cuproptosis-related genes were obtained from previous literature^19^. Ferroptosis-related genes were downloaded from Ferrdb^20^. A list of 606 nonapoptotic-RCD-related genes (NRGs) are shown in **Table S1**.

Kaplan-Meier survival analysis (p<0.05) was used to screen 5756 genes at mRNA level and 1967 genes at protein level with prognostic value based on the expression matrix from FU-iCCA cohort.

### Somatic mutation and copy number alteration analysis

DNA alteration includes mutation, which means truncating and missense, and copy number alteration. For TCGA-CHOL, we have downloaded the RNA-seq data in fragments per kilobase of exon per million mapped fragments (FPKM) values and matched clinical data from the UCSC Xena data portal. Somatic mutation and CNV data were obtained via using the R package TCGAbiolinks ^21^.We have analyzed the information about Somatic mutation data which was downloaded in the form of Mutation Annotation Format (maf) and then used them to calculate TMB using the R package maftools ^22^.

### Consensus Clustering

The number of NRG-related clusters in FU-iCCA cohort and their stability were defined by the consensus clustering algorithm which have used the R package ‘ConsensusClusterPlus’ with 1,000 repetitions ^23^. Principal component analysis (PCA) was performed to show the distribution difference of NRG-related clusters. For the purpose of evaluating the clinical value of the NRG-related clusters, we have explored the relationships between NRG-related clusters with prognosis and other clinicopathological features such as adjuvant therapies, glutamyltransferase, ALT aminoleucine transferase, ALB albumin, TB total bilirubin, distal metastasis, regional lymph node metastasis, liver cirrhosis, vascular invasion, biliary tract stone disease, liver fluke, intrahepatic metastasis, age, sex tumor size diameter, TNM stage, perineural invasion, preoperative serum AFP, CA19 carbohydrate antigen, and CEA carcinoembryonic antigen.

### Univariate and Multivariable Regression

We performed univariate Cox regression on FU-iCCA with gene expression and overall survival. Multivariate Cox regression was used to evaluate independent risk factors in the same cohort. And genes and factors with a false discovery rate (FDR) <0.05 were considered statistically which is associated with patient survival. The results of univariate and multivariate Cox regression were acquired and visualized by using the R package forest plot.

### Creating and confirming the predictive NRG risk score

By computing the overall value of risk, the value of NRG was established. And then the train sets were utilized to generate an NRG risk score for prognosis. All patients in FU-iCCA cohort were divided into train (n=144) and test (n=63) sets. In order to minimize the possibility of overfitting the model, the glmnet package in R was used to perform least-squares regressions and selection operator regressions. Candidate genes were selected via using multivariate Cox analysis to establish a prognostic NRG risk score in the training set. Both the train set and the test set were splited into groups of high-risk and low-risk based on their risk ratings. In each set, Kaplan-Meier analyses of survival and ROC curves were conducted.

### Creation and verification of a nomogram

A nomogram was created by using the rms program to predict overall survival, which is based on clinically significant characteristics and the NRG risk score. Each clinicopathologically significant characteristic was given a score by the means of the nomogram model, and the overall score was obtained by summing all the individual scores. By contrasting the area under the time-dependent ROC curves of survival rates after one, two, and three years, the nomogram’s accuracy in predicting survival rates was also validated. Additionally, model calibration was performed for the purpose of comparing the predicted likelihood of survival outcomes across the 1-, 2-, and 3-year periods with the actual survival occurrences.

### Identification of Differentially Expressed Genes (DEGs) and Functional Enrichment Analysis

DEGs between two NRG-related clusters and between high- and low-risk groups were determined based on t tests by using the R package ‘limma’ from proteome (protein), transcriptome (mRNA) and phosphoproteome respectively. An adjusted P value <0.05 was considered statistically significant. These significant genes which were then performed KEGG pathway or GO enrichment analysis by using clusterProfiler R package and an FDR value <0.05 or other thresholds were considered as the cutoffs of significantly regulated pathways.

### Correlations of molecular subtypes or risk score with TME, PD-1, and PD-L1 in iCCA

For the purpose of calculating the total number of tumor-infiltrating immune cells and subgroups of immune cells in each sample, we have used the ESTIMATE algorithm to evaluate the immune and stromal scores of each patient between the cluster 1 and cluster 2 or the high-risk and low-risk groups. In addition, the fractions of 23 human immune cell subsets of every iCCA sample were calculated by the ESTIMATE algorithm. Furthermore, the levels of immune cell infiltration in the iCCA TME were also determined by using a single-sample gene set enrichment analysis (ssGSEA) algorithm. This was done to assess how many immune cells had invaded the tumor altogether. We also analyzed the correlations between the two subtypes of PD-1 and PD-L1 expression. The IMvigor210 dataset included a total of 298 iCCA cases with expression data, complete survival data, follow-up information and immune therapy effect information which were downloaded from the IMvigor210CoreBiologies R package. GSE78220 included 27 iCCA patients with entire clinical data and the efficacy of immune therapy. According to the patients’ response to immune therapy, patients in IMvigor210 and GSE78220 cohorts were sorted into four groups: progressive disease (PD), stable disease (SD), partial response (PR) and complete response (CR).

### scRNA-Seq Data Analysis

The quality control (QC) and cell clustering have been analysed on the integrated dataset which is based on t-SNE algorithm implemented in Seurat following the online pipeline (https://satijalab.org/seurat/). CellChat (http://www.cellchat.org/) was used to analyze the intercellular communication networks from scRNA-seq data.

### Analysis of drug susceptibility

For the purpose of exploring differences in the therapeutic effects of chemotherapeutic drugs in patients between two NRG-related clusters, and high- and low-risk groups, we have calculated the semi-inhibitory concentration (IC50) values of chemotherapeutic drugs commonly which is used to treat iCCA by using the “pRRophetic” package.

### Proteome and phosphoproteome data analysis

A total of 8,320 proteins and 18,347 phosphosites which were quantified from the FU-iCCA cohort ^17^. In order to obtain the protein level phosphorylation data, to begin with, we collapsed the phosphorylation sites from the 10 NRG-related proteins with the median ratio and then divided iCCA samples into high- and low-phosphorylation groups. Then combine the protein expression matrix from FU-iCCA cohort to analyze the NRG expression level in the two subgroups.

### Tissue samples

Five pairs iCCA and nearby non-tumor tissues were harvested from iCCA patients at the Shandong Provincial Hospital Affiliated to Shandong University. The samples were preserved at −80°C till use. All the individuals in this study have offered their written informed consent. The study was permitted by the Ethics Committee of Shandong Provincial Hospital Affiliated to Shandong University.

### Western blotting analysis

Total protein from tissues were extracted using standard methods and protein concentrations were determined by a BCA protein assay kit. Equal of protein was separated by 10% SDS-PAGE gel and transferred to a PVDF membrane. The membrane was blocked and shaken in 5% skim milk and then incubated with the primary antibodies, including anti-VPS37B (1:500; Proteintech, 15653-1-AP), anti-CHMP2B (1:1000;Abcam, ab157208), anti-CASP3 (1:500; Abcam, ab13847), anti-CP (1:500; Abcam, ab48614), anti-CIRBP (1:1000; CST, #68522S), anti-CYB5R1 (1:500; Invitrogen, PA5-21506), anti-PRKACA (1:1000; Abcam, ab32376), anti-NF2 (1:5000; Abcam, ab109244), anti-GPX4 (1:2000; Proteintech, 14432-1-AP), anti-HRAS (1:500; Proteintech, 18295-1-AP), anti-TGF-β (1:1000; Proteintech, 21898-1-AP) and anti-GAPDH (1:3000; Santa Cruz, sc-47724), and the corresponding secondary antibodies for protein detection. Then, we used the fully automatic chemiluminescence/fluorescence image analysis system (Tanon 5200, Shanghai, China) to observe the bands with AllDOC-X software (Exposure time; 0.05s 0.5s 1s 5s 10s). The Image J (1.51j8) gel analysis software was utilized to quantitatively analyze the band intensities, normalizing with GAPDH.

## Statistical analyses

All statistical analyses were performed which used R version 4.1.0. Statistical significance was set at *p* < 0.05.

## Results

### Identification of nonapoptotic-RCD-related genes (NRGs) from iCCA through the transcription and protein levels

To fully understand the expression pattern of NRGs involved in tumorigenesis, the information of 207 patients from FU-iCCA cohort and previous reports were used in our study for further analysis. Detailed information on the 207 iCCA patients is presented in **Table S2**. OS evaluation of all genes in the FU-iCCA expression spectrum matrix was performed by Kaplan-Meier analysis (p<0.05) which has brought about 5756 OS-associated genes at mRNA level and 1967 OS-associated genes at protein level. These OS-associated genes from transcription and protein levels and 606 NRGs were then intersected to obtain 52 NRGs with the prognostic values both in mRNA and protein levels in patients with iCCA (Figure 2A). The comprehensive landscape of interactions, regulator connections of these 52 NRGs and their degree of interaction was demonstrated in network (Figure 2B). To further explore the expression characteristics of NRGs in iCCA, we have used a consensus clustering algorithm to categorize the patients with ICC based on the expression profiles of the 52 NRGs. Our results showed that k = 2 appeared to be an optimal selection for sorting the entire cohort into cluster 1 (n = 68) and 2 (n = 139) (Figure 2C). The significant differences in NRGs profiles between the two clusters have been revealed via the PCA analysis (Figure 2D). The result which means a longer OS in patients with cluster 2 than that in patients with cluster 1 has been shown in the Kaplan-Meier curves (*p* <0.0001; Figure 2E). Furthermore, the significant differences in 52 NRGs’ expression and 21 clinicopathological characteristics can be revealed by the comparisons of the clinicopathological features of the two clusters of iCCA (Figure 2F). As shown in Figure 2F, compared with cluser 1, most NRGs of cluster 2 at mRNA level were preferentially related to lower tumor size diameter, lower TNM stage, less perineural invasion, with preoperative serum AFP, lower CA19 carbohydrate antigen, and lower CEA carcinoembryonic antigen. Conversely, most NRGs had the opposite relationship at the protein level with the above clinical features. These indicated that NRGs may affect iCCA development by some means of the potential mechanisms, which shows the differences at the transcriptional and protein levels.

**Figure 2.**
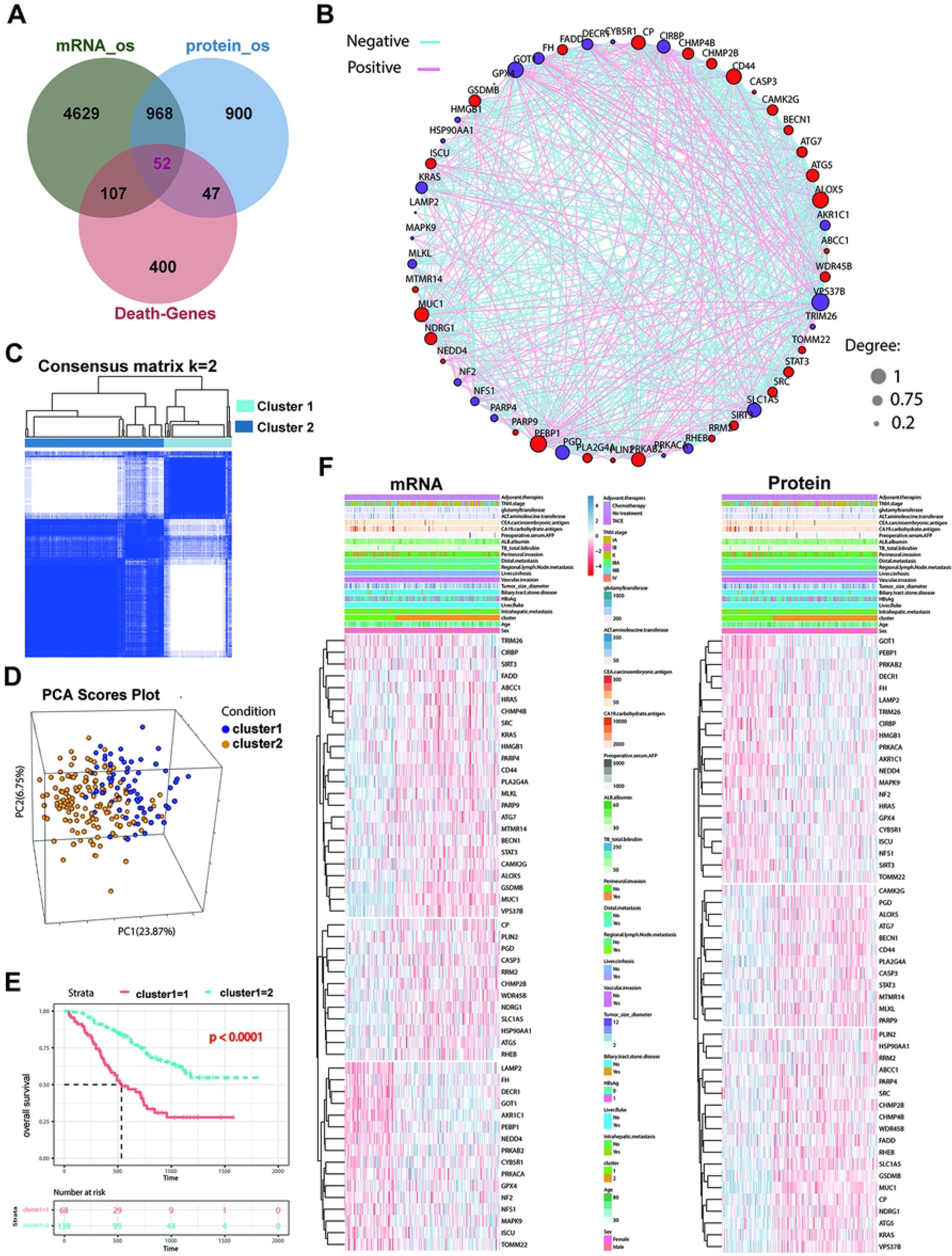
Identification of nonapoptotic-RCD-related genes (NRGs) from iCCA through the transcription and protein levels. (A) Venn diagram of mRNAs/proteins having significant prognostic values based on all genes from FU-iCCA expression spectrum matrix and 606 nonapoptotic-RCD-related genes (NRGs). (B) Interactions of 52 NRGs. The line connecting the NRGs represents their interaction, with circle size indicating the degree of the association between NRGs. Green and pink represent negative and pink positive correlations, respectively. (C) Consensus matrix heatmap defining two clusters (k = 2) and their correlation area. (D) PCA analysis showing a remarkable difference in transcriptomes between the two clusters. (E) Cluster-specific Kaplan-Meier OS curves based on their transcriptomes in the FU-iCCA cohort. (F) NRG expression at mRNA and protein levels, and clinicopathological traits vary across two clusters.

### Construction and validation of the NRG-related prognostic signature (NPS) model

The NPS model was established based on 52 NRGs. To begin with, we used the “caret package” in R to randomly classify the patients into training (n = 144) and testing (n = 63) cohorts. LASSO analyses for 52 NRGs were performed to further select optimum prognostic signature. At the minimum of λ value, 10 core NRGs were selected to construct NPS model (**Figs.3A**), and their LASSO coefficient were shown in **Table. S3**. The NPS risk score was constructed as follows:

Risk score = (0.411744* expression of VPS37B) + (−0.38376* expression of CIRBP) + (0.357305* expression of CHMP2B) + (−0.32076* expression of CYB5R1) + (-0.31491* expression of PRKACA) + (-0.31118* expression of NF2) + (0.298858* expression of CASP3) + (-0.22188*expression of GPX4) + (-0.20487*expression of HRAS) + (0.192705* expression of CP).

According to the medium vale of the risk scores, patients with iCCA in the training set were sorted into the groups with low- and high-NPS risk score. The iCCA patients with high-NPS risk score high risk had poor OS (p<0.0001; Fig. 3B**, left**) with 1-, 2- and 3-year AUC values were 0.78, 0.82 and 0.8 (Fig. 3B, **right**). In testing set, high risk group had poor OS (p=0.02; Fig. 3C**, right**) with 1-, 2- and 3-year AUC values were 0.76, 0.68 and 0.85 (Fig. 3C**, right**). We then performed multivariate Cox regression analysis to finally obtain four high-risk genes (*VPS37B, CHMP2B, CASP3, and CP*) and six low-risk genes (*CIRBP, CYB5R1, PRKACA, NF2, GPX4, and HRAS*) in training set (Fig. 3D**, upper**). In testing set, the forest plot indicates that (*VPS37B, CHMP2B, NF2, CASP3, and CP* were risk factors with Hazard ratio (HR) >1, while *CIRBP, CYB5R1, PRKACA, GPX4, and HRAS* were protective factors with HR<1 for iCCA patients (Fig. 3D**, bottom**). Considered the inconvenience clinical utility of NPS-risk score in predicting OS in patients with iCCA, a nomogram which has incorporated the NPS risk score and 12 clinicopathological parameters was established to predict the 1-, 2-, and 3-year OS rates (**Figure S1A**). Each patient’s total point values were determined based on prognostic characteristics such as their age, level of risk score, and the TNM stage included. The subsequent calibration plots showed that the proposed nomogram performed better than an ideal model (**Figure S1B).** Furthermore, we also compared the response of patients between different clusters and between different risk score groups to the 21 clinical characteristics from Figure 1F, and found that the proportion of patients with cluster 1/2 (**Figure S2A**) and high-/low-NPS risk scores (**Figure S2B**) had expressed significant difference for 12 clinical characteristics, as well as the trend of the proportion of patients in the two classification methods is consistent.

**Figure 3.**
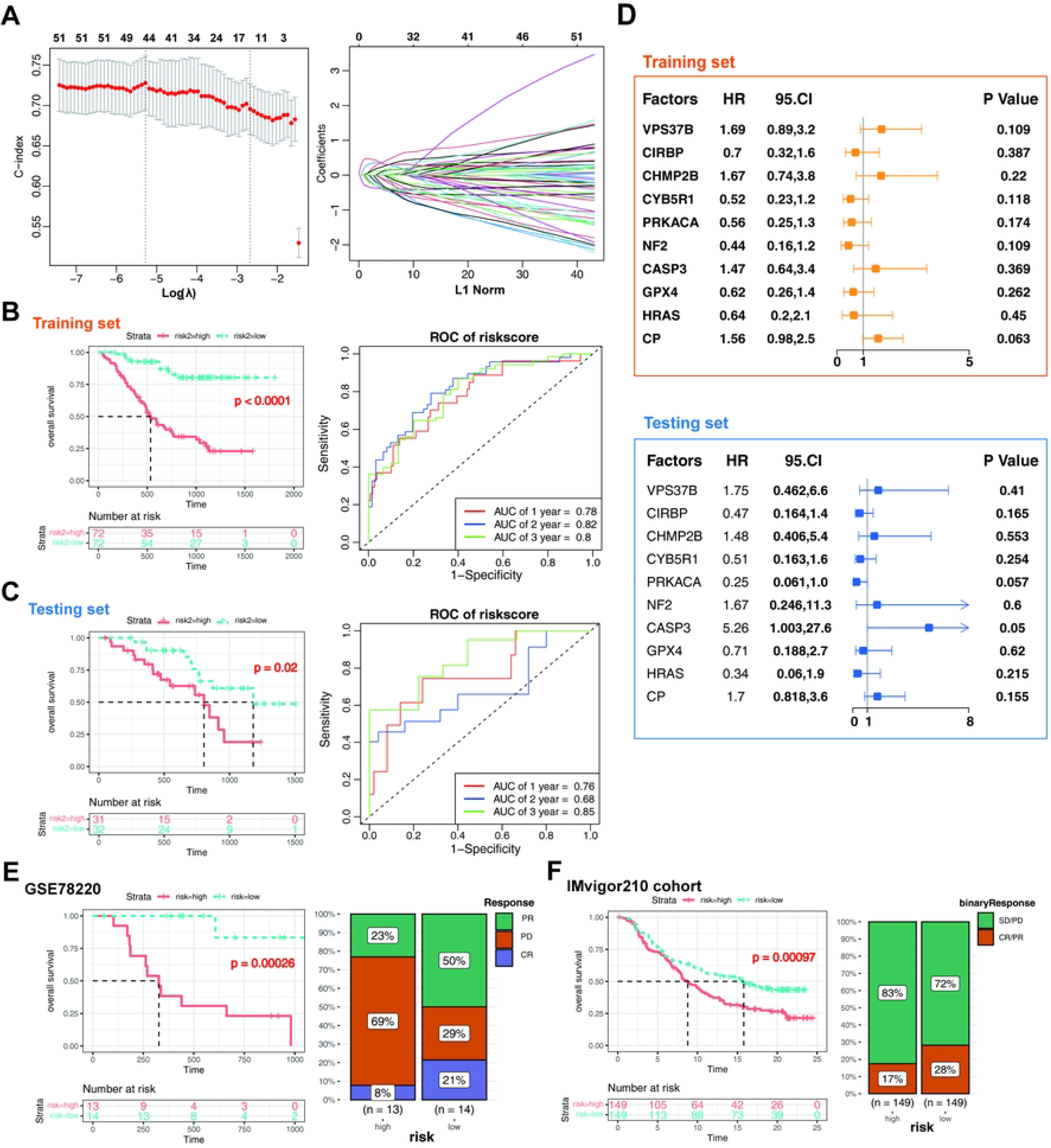
Construction and validation of the NRG-related prognostic signature (NPS) model. (A) The LASSO regression analysis and cross-validation of LASSO regression parameter selection based on 52 NRGs. (B-C) Kaplan-Meier analysis of the overall survival between the high- and low-NRG-risk score groups and ROC curves for forecasting 1-, 2-, and 3-year OS in the (B) training set and (C) testing set, respectively. (D) Forest plot of multivariate cox regression analysis for 10 core prognostic NRGs in the training set and testing set, respectively. (E-F) Survival analysis for patients with high- and low-NPS-risk score in the GSE78220 melanoma cohort and IMvigor210 cohort (left). Bar graph illustrated the treatment response [complete response (CR)/partial response (PR) and stable disease (SD)/progressive disease (PD)] to immunotherapy in high and low-NPS risk score subgroups in GSE78220 and IMvigor210cohorts (right).

To further investigate whether NPS risk score could predict patients’ response to ICI therapy in the immunotherapy cohort, we were going to explore the prognostic value of NPS-risk score on patients who received ICI therapy in the GSE78220 and IMvigor210 cohorts by dividing them into high- and low-NPS risk score groups. Patients with a higher-NPS risk score had significantly shorter overall survival than those with a lower NPS risk score in both the GSE78220 and IMvigor210 cohorts (**Figs. 3E-F, left**). Meanwhile, the proportion of patients with low-NPS-risk score who responded to immune checkpoint blockade therapy (PR/CR) was larger than the proportion of patients with high-NPS-risk score (SD/PD) both in the GSE78220 and IMvigor210 cohorts (**Figs. 3E-F, right**). The subsequent forest plot indicates that (*PRKACA, CYB5R1, CHMP2B, and CASP3* were risk factors with Hazard ratio (HR) >1, while *VPS37B, NF2, CIRBP, GPX4, and CP* were protective factors with HR<1 based on the GSE78220 melanoma cohort (**Figure S1C**). However, *CP* was a risk factor with Hazard ratio (HR) >1 in the IMvigor210 cohort (**Figure S1D)**. In a nutshell, these findings suggest that the NPS-risk score may be a novel indicator of the response of iCCA to ICI therapy.

### NRGs in iCCA: Expression, genetic variants, and prognostic values

Next, we explored somatic copy number alterations in these NRGs and discovered prevalent copy number alterations in all 10 NRGs. Among them, the copy number variation (CNV) and VPS37B have generally increased, while CIRBPCP, *CYB5R1, GPX4, and HRAS* showed CNV decreases (Figure 4A). Figure 4B shows the locations of the CNV alterations of 10 NRGs on their respective chromosomes. It is worth noting that we have found that GPX4, CIRBP and PRKACA shared similar mutation frequencies and similar patterns of CNV (Figure 4B). We divided 36 CHOL samples from TCGA into high and low risk groups, and a lower level of risk score is linked to longer OS (Fig. 4C). We then analyzed the distribution variations of the somatic mutations between two groups in the TCGA-CHOL cohort. The top five mutant genes in the high-risk group were *PBRM1, ARID1A, BAP1, EPHA2* and *ARAF*, and the top five mutant genes in the low-risk group were *IDH1, ARID1A, CHD7, EPHA2* and *FBN3* (Fig. 4D). Patients with a high- or low-risk score all had frequencies of *ARID1A* and *EPHA2* mutations, indicating that these two gene mutations are closely related to CHOL progression. On the FU-iCCA cohort, there were 10 NRGs from 207 iCCA patients’ expression levels examined in the NPS model were examined in different clusters or in high-/low- risk score groups. Significant differences in their expressions can be observed not only between the cluster1 and cluster2 (Fig. 4E) but also between high-NPS risk score and low-NPS risk score groups (Fig. 4F). In conclusion, these evidences indicated that the expression and mutation patterns of NRGs are highly heterogeneous in iCCA, which further suggested that CNV alterations might have the ability to modify the way NRGs are expressed.

**Figure 4.**
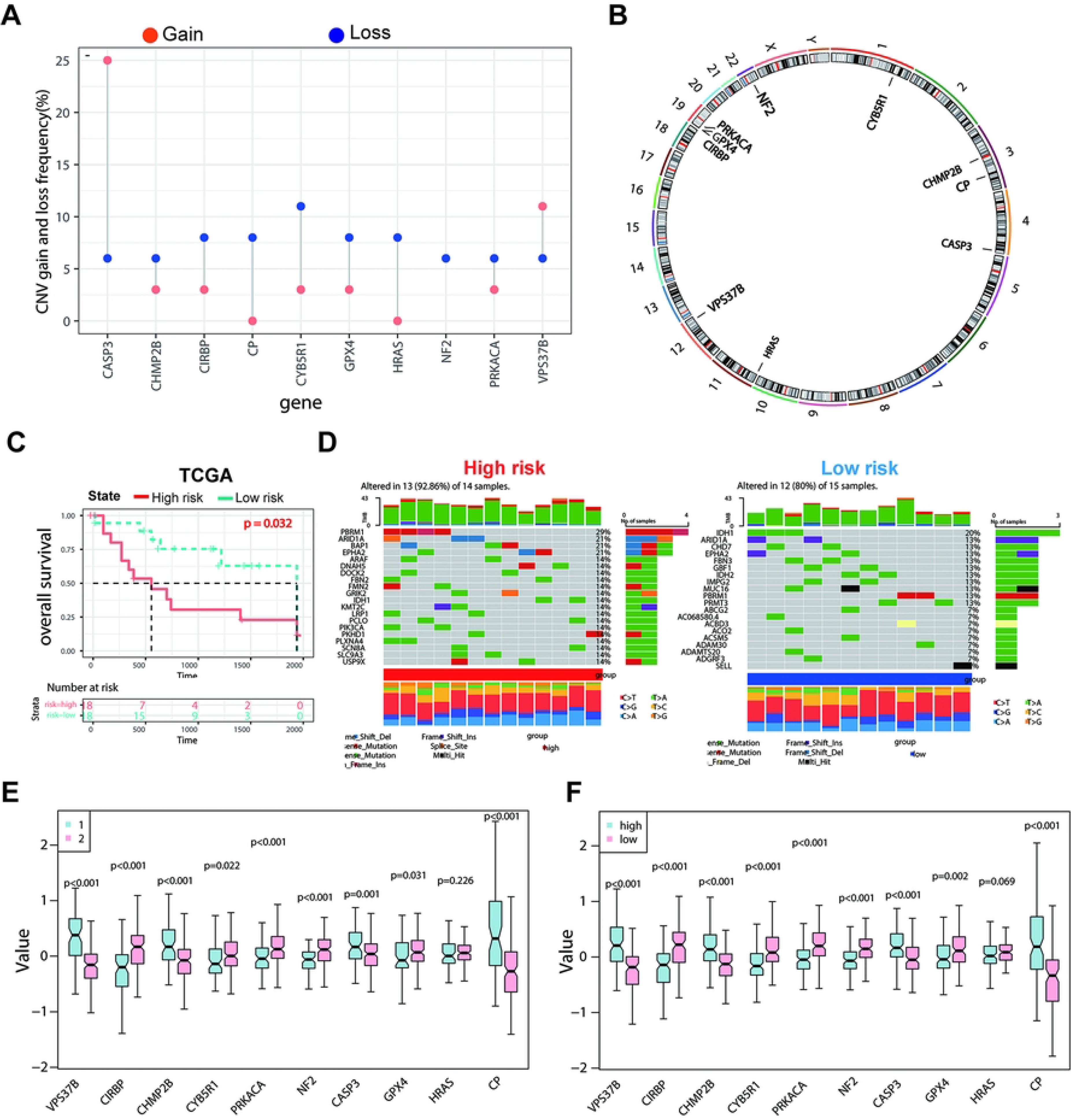
Landscape of genetic and expression variation of 10 NRGs, and association in the TCGA-CHOL cohort. (A) The frequency of CNV gain and loss, and non-CNV of 10 NRGs. (B) The sites of CNV variation in CRGs on the 23 chromosomes. (C) Kaplan-Meier curves showing that the association between NPS-risk score and overall survival in the TCGA-CHOL cohort (n=36). (D) The waterfall plot of somatic mutation features established with high and low NPS risk scores. Each column represented an individual patient. The upper barplot showed TMB, the number on the right indicated the mutation frequency in each gene. The right barplot showed the proportion of each variant type. (E) The expression of the 10 NRGs in iCCA patients between cluster 1 and cluster 2. (F) The expression of the 10 NRGs in iCCA patients between high- and low-NPS-risk score groups.

### Analyses of TME infiltration and functional enrichment in distinct clusters and high-/low- NPS risk score groups

To explore the differences in underlying biological function among the two NRG-related clusters, we performed GSVA on the two clusters. Cluster 1 showed enrichment in GO terms which is associated with negative regulation of transcription by competitive promoter binding, positive regulation of transcription from RNA polymerase Ⅱ promoter in response to stress, regulation of voltage gated sobium channel activity and secretion by tissue etc. (**Figure S3A**). We also applied GSVA enrichment analysis to observe the possible effects of the two clusters on biological behavior. Compared with cluster 2, cluster 1 had an enrichment in the pathways linked to glycan biosynthesis, metabolic-activated and immune activation pathways, such as the glycan biosynthesis, mTOR signaling pathway, glycerophospholipid metabolism, MAPK signaling pathway, B cell receptor signaling pathway, nature killer cell mediated cytotoxicity etc. (**Figure S3B**).

By comparing the differential functional enrichment of multi-omics data between two clusters, we observed that cluster 1 showed upregulation of hypoxia, apoptosis, epithelial mesenchymal transition (EMT), and glycolysis (Fig. 5A). For the purpose of exploring the differences in underlying biological function among the high- and low-NPS risk score, we have performed GSVA on the two groups. Interestingly, while performing the same analysis of the iCCA samples of the high- and low-NPS risk groups as those from Cluster 1 and Cluster 2, all the characteristics of cluster 1 were found to be in the accordance with the high-risk group (**Figure S3C-D**). The pathway enrichment of the RNA, protein and phosphorylation levels indicated that the group with high-NPS risk score was also mainly enriched in the hypoxia and glycolysis (Fig. 5B). Furthermore, the representative proteins of the EMT process (FN1, COL7A1, ECM1, and LAMC1) and apoptosis pathways (CASP3, BCL10, and ERBB2) were significantly overexpressed in the cluster 1. However, EMT-related SSP1 and apoptosis-related KRT18 were obviously overexpressed in the cluster 2 (Fig.5C). Besides, the representative proteins of hypoxia (ALDOA, NDRG1, ANXA2, and CP) and glycolysis (PGK1, LDHA, MET, and PKP2) were significantly overexpressed in the cluster 1 and high-NPS risk score groups (Fig. 5C**-D**). Then, the pathway and process enrichment analysis based on the differentially phosphorylated sites were performed as well. We found that many pathways and functions were significantly enriched, such as mRNA metabolic process, neutrophil degranulation, VEGFA-VEGFR2 signaling pathway, and signaling by Rho GTPases, and that most of these functions and pathways were mediated by cluster 1 and high-NPS risk score groups (Fig. 5E**-F**).

**Figure 5.**
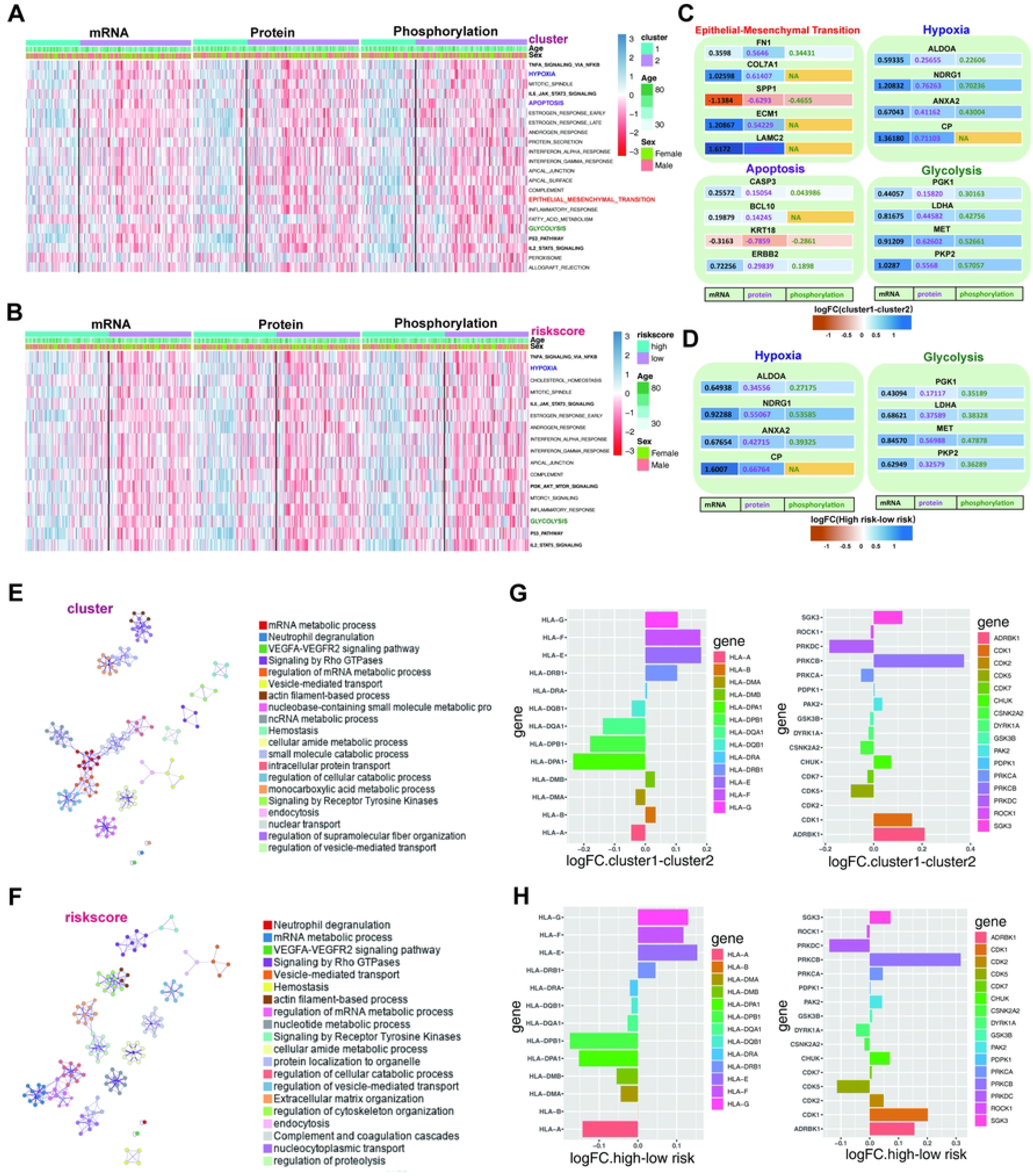
Analyses of functional enrichment. (A) Integrated analysis of altered pathways at transcriptome, proteome, and phosphosproteome levels among cluster 1 and cluster 2. (B) Heatmap shows the representative biological pathways among high-NPS risk score and low-NPS risk score subgroups in RNA, protein, phosphoprotein levels of FU-iCCA cohort. (C) The differential expression level of representative proteins of EMT, apoptosis, hypoxia and glycolysis between cluster 1 and cluster 2. The color represents the logFC (cluster1-cluster2) ratio of protein expression in RNA, protein, phosphoprotein levels, respectively. (D) The differential expression level of representative proteins of hypoxia and glycolysis between high- and low-NPS risk score subgroups. The color represents the logFC (high risk-low risk score) ratio of protein expression in RNA, protein, phosphoprotein levels, respectively. (E) The expression of phosphorylated protein sites in cluster 1 and cluster 2 subgroups were applied to perform pathway and process enrichment analysis via Metascape. (F) The expression of phosphorylated protein sites in high- and low-NPS risk score subgroups were applied to perform pathway and process enrichment analysis via Metascape. (G) Gene expression of HLA and kinases gene sets between cluster 1 and cluster 2 subgroups. (H) Gene expression of HLA and kinases gene sets between high- and low-NPS risk score subgroups.

The results also suggested that DEGs which were highly expressed in high-NPS risk score significantly enriched in the B-cell receptor signaling pathway, T cell receptor signaling pathway, nature killer cell mediated cytotoxicity mTOR signaling pathway, acute myeloid leukemia, VEGF signaling pathway, p53 signaling pathway, and MAPK signaling pathway (**Figure S3B**). Considering the close relationship between distinct clusters/risk score and immune activity, we have used the ImmuneCellAI online tool to study the TME of two clusters or high-/low- risk score groups based on the TCGA cohort (**Figure S3E-F)**. The cluster 1 subtype was characterized by the high infiltration of DC (dendritic cell), eosinophils, epithelial cells, erythrocytes, keratinocytes, macrophages, monocytes, neutrophils etc., whereas the cluster 2 was featured by the high infiltration of B cell, CD4 memory T cells, CD4 + T cells, CD8 native T cells, CD8+ T cells, Th1 cells, Tregs, etc. (**Figure S3E**). Similarly, the high- NPS risk score subgroup was characterized by the high infiltration of DC (dendritic cell), eosinophils, epithelial cells, erythrocytes, keratinocytes, macrophages, monocytes, neutrophils sebocytes, etc., while the low-NPS risk score subgroup was featured by the high infiltration of B cell, CD4 memory T cells, CD4 + T cells, CD8 native T cells, CD8+ T cells, MEP, myocytes, NK cells, Th1 cells, Tregs, etc. (**Figure S3F**). The Stromalscore, ImmuneScore and ESTIMATEScore in CHOL tumor tissues were calculated using the ESTIMATE algorithm based on the TCGA expression profiles. ESTIMATE generates a stromal score which has the ability to meansure the presence of tumor-associated stroma and an immune score that represents the infiltration level of the immune cells and combines them to produce an index termed ‘estimatescore’ that comprehensively infers tumor purity. As shown in **Figure S3G-H**, compared with those of cluster 2 and low-NPS risk score subgroups (*P*<0.05), samples in cluster 1 and high-NPS risk score also exhibited significantly higher than ESTIMATEScore and ImmuneScore. Then, we investigated the relationship of two subtypes with human leukocyte antigen (HLA) stimulators and protein kinases. There has been a higher expression lever of HLA-B, HLA-DMB and HLA-E-G in cluster 1 and high-NPS risk score subgroups, while HLA-A, HLA-DPA1, DPB1, DQA1, and DQB1 obtain the higher expression level in the cluster 2 and low-NPS risk score subgroups (Fig.5G **and 5H**). Through the further enrichment analysis of the TCGA-CHOL cohorts, the significant enrichment of immune-related kinases PRKCB, cyclin-dependent kinases CDK1/2, and the apoptosis-related kinases PAK2 and SGK3 were revealed in the cluster 1 and high-NPS risk score subgroups; however, PRKDC, CSNK2A2, and CDK5 were markedly enriched in the cluster 2 and low-NPS risk score subgroups (**Fig5G-5H**). These findings indicated that the clustering and scoring models in this study were matched.

### High NRG risk score implies an immune-active tumor microenvironment

Next, we sought to identify key players in the TME contributing to NRG-related phenotypes. And we obtained Single-cell mRNA profiles of seven primary tumor and one normal tissue sample by means of the GSE151530 dataset. After quality control and removal of batch effects, filtered cells were clustered and annotated into 6 major cell types, including CD8+ T cells, erythroid cells, HSCs, monocytes, B cells and CD4+ T cells (Fig. 5B**-C**), of which only four kinds showed significant differences in cell infiltration levels between high- and low-NPS risk score subgroups (Fig. 6A). Compared with low-NPS risk score subgroup, the proportion of CD4+ T cells, HSCs and erythroid cells was also higher in the high-NPS risk score subgroup (Fig. 6D). For the purpose of characterizing intercellular interactions in high- and low-risk group, we applied the CellChat to infer putative cell-to-cell interactions of 6 major cell types based on ligand-receptor signaling. Interestingly, the enhencement intercellular interactions for the low-risk group could be observed (Fig. 6E), where CD4+ T cells displayed widespread communication with other cell types, indicating that they were mainly potential contributors to the NPS-related phenotype. We then dissected the signaling networks in order to identify individual ligand-receptor pairs which were featured in the high- and low-risk samples. Significant differences in the patterns and intensity of multiple cellular communications can be observed between the high- and low-risk groups (Fig. 6F). Such as, enhanced signaling from CD4+ T cells to erythroid cells and CD8+ T cells were observed in the high-NPS risk score subgroup, including VEGFB-VEGFR1, VEGFA-VEGFR, MDK-SDC1/2/4. From a detailed analysis of the cell- chat which brought about the list of all igand-receptor pairs of intercellular communication, the increased communication probability between CD4+ T cells and other five major immune cell types via MIF-(CD74 + CXCR4), MIF-(CD74 + CD44), MDK-NCL, and the enhanced recruitment of immune cells by CD4+ T cells in the NPS risk score high group were found. Herein, the hypothesis of potential mechanism of between them was demonstrated in Fig. 6G. Different NPS risk score subgroup might have different communication strengths and ligand-receptor pairs suggesting that NPS-related TME cells might have more interactions with tumor cells and thus contribute to the progress of iCCA. Collectively, these results have confirmed the positive relation between the NPS and the immune activity in iCCA.

**Figure 6.**
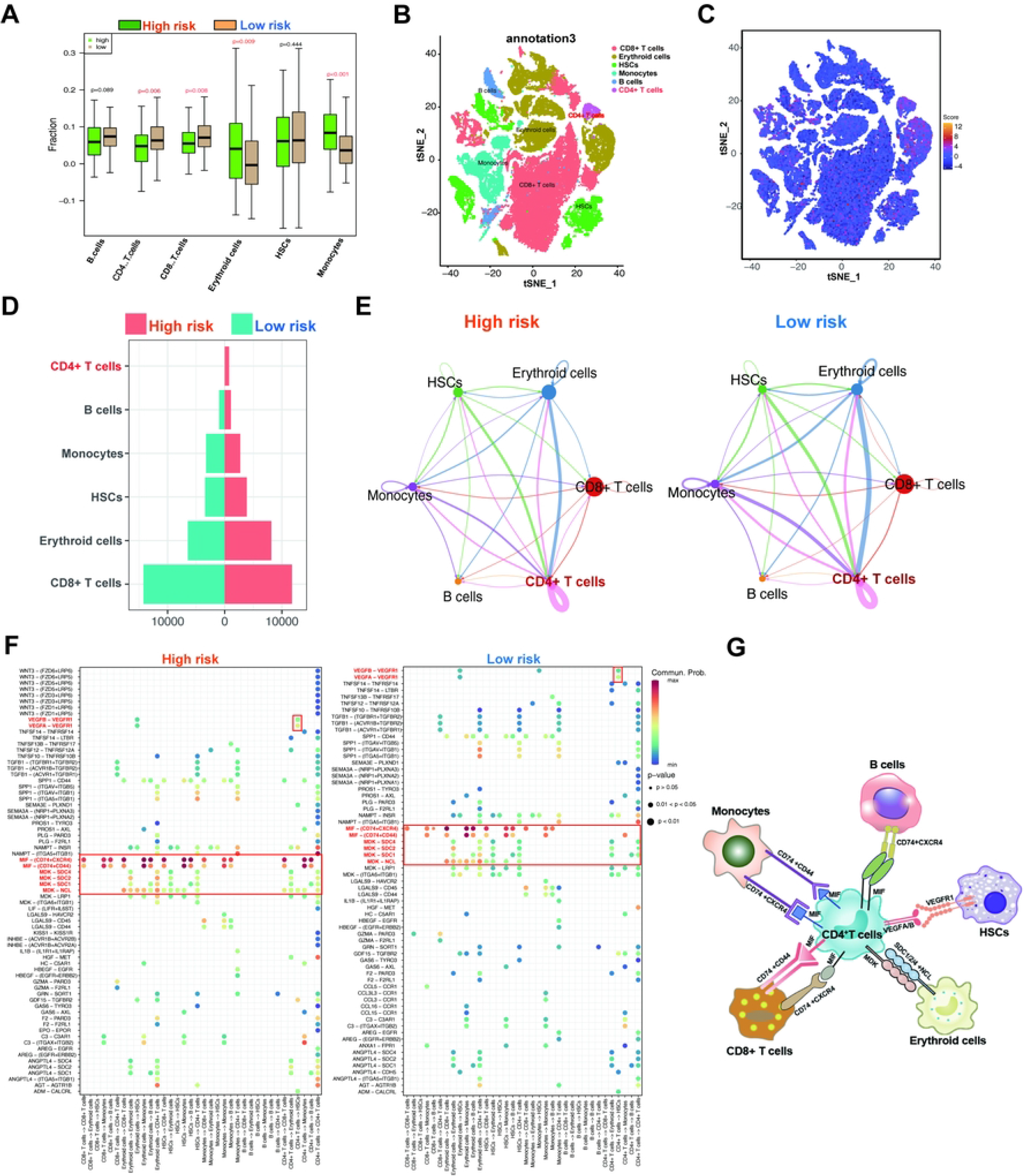
High NPS risk score implies an immune-active tumor microenvironment in the single cell cohort. (A) The abundance of six TME-infiltrating cell type in high- and low-NPS risk score subgroups. (B) t-SNE plot showing the composition of 6 main cell subtypes derived from iCCA samples. (C) The dynamics of NPS risk score in 6 main cell types are represented in the t-SNE plot. (D) The proportion of 6 main cell types in the high or low NPS risk score subgroups. (E) Cell-Cell communications between main six cell types from high-risk (left) and low-risk (right) group by Cell chat analysis. The brand links pairs of interacting cell types. (F) The significantly related ligand-receptor interactions from main six cell types between high- and low-NPS risk score subgroups. (G) Hypothesis of the mechanism of NPS risk score-related TME cells affecting cell communication.

### scRNA-seq Reveals Transcriptional Profiles and cell communication pattern of CD4+ T cells in iCCA

In view of the above results suggesting a potential role of CD4+T cells in the iCCA tumor microenvironment, we analyzed the transcriptional profile of CD4+T cells. There were 10 NRGs in the NPS model expressing differently in 6 major cell types (Fig. 7A **and S4**). It is worth noting that all NRGs were expressed most strongly in the CD4+ T cells, of which CP was the highest expressed (Fig.7B). Interestingly, compared to the low-NPS risk score subgroup based on the scRNA-seq data, CP was also significantly overexpressed in CD4+ T cells in the high-NPS risk score subgroup (Fig.7C). According to the results of cell communication (Figure 5E), the closest communication between CD4+ T cells and erythroid cells can be found, while we can also find that top enriched TGF-β signaling network was strengthened, which was between CD4+ T cells and erythroid cells is significantly different in the high- and low-NPS risk score groups, suggesting that TGF-β signaling pathway play a crucial role in the progression of iCCA (Fig.7D). We then found that there was TGF-β widely expressed in six cell types, which is mainly concentrated in CD8+T cells and erythroid cell with an interesting phenomenon, which was observed that TGF-β was only highly expressed in one part of erythroid cells, while its expression was lower in the other part (Fig.7E). Moreover, these two erythroid cells subclusters were clearly separated,based on this feature, the erythroid cells were annotated into TGF-β positive (+) and negative (-) subclusters (Fig.7G). We further focused on the cell communication between these two erythroid cells subclusters and CD4+T cells, and the enrichment of CP in CD4+T cells which interact with TGF-β from the erythroid cells-TGF-β+ were found (Fig.7F). We then analyzed the communication between erythroid cells-TGF-β+ and other five types of cells, of which results confirmed that erythroid cells-TGF-β+ and CD4+ cells communicate at high frequency through the TGF-b signaling pathway (Fig.7H**-I**). We suspect that erythroid cells-TGF-β+ and CD4+T cells may communicate in the way of the secretion of CP and TGF-b proteins which are highly abundant in them (Fig.7J). The findings of a new molecular feature in the progression of iCCA were revealed.

**Figure 7.**
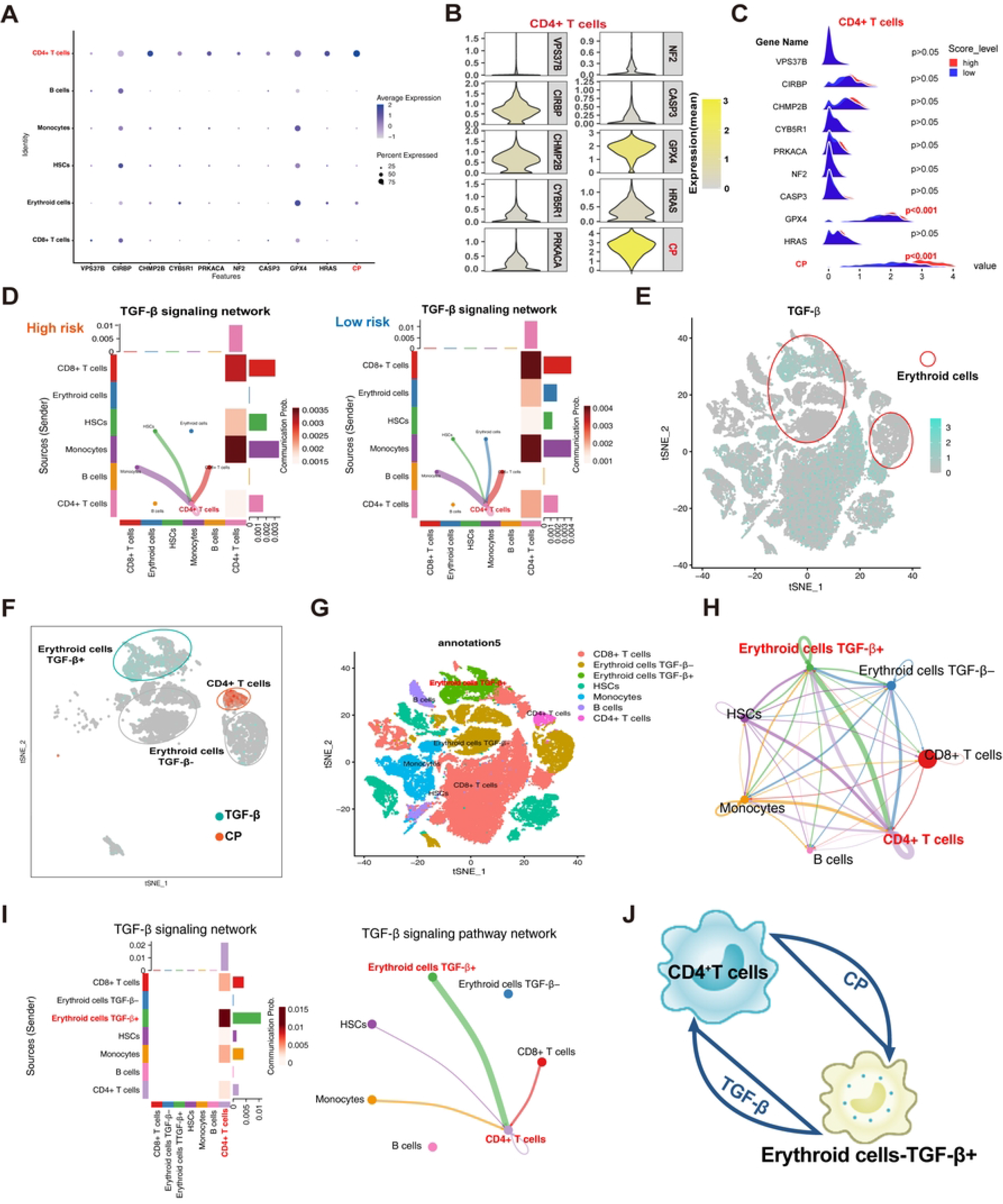
scRNA-seq Reveals Transcriptional Profiles and cell communication pattern of CD4+ T cells in iCCA. (A) Dot plot showing the expression of 10 NRGs in 6 main cell subtypes. Dot size represents the percent of expressing an NRG in a particular cell type and intensity of color indicates the average expression level in that cell type. (B) Violin plots showing the expression of 10 NRGs enriched in CD4+ T cells. (C) The peak maps showing the expression of 10 NRGs among high- and low-NPS risk score subgroups. (D) Circos plots showing cell–cell communications from main six cell types of the TGF-β signaling pathways between high-risk (left) and low-risk (right) subgroup by Cell chat analysis. Bar plot showing the number and percentage of main six cell types. (E) Distribution of TGF-β in all six immune cell types overlaid on the 2D t-SNE plot. (F) Distribution of the indicated genes (*CP* and *TGF-β*) in CD4+T cells and erythroid cells overlaid on the 2D t-SNE plot. (G) Cell type annotations by using the Seurat t-distributed stochastic neighbor embedding (t-SNE) plot. (H) Interaction among different cell types by Cell chat analysis. The width of links represents the number of significant ligand-receptor interactions between the indicated cell types. (I) Cell-Cell communications from CD4+T cells to other cells based on the TGF-β signaling network. (J) Schematic representation of CD4+T cells interaction with erythroid cells-TGF-β+.

### Identification of NPS risk score-related drugs for iCCA

Then, we selected the current chemotherapy drugs used for the treatment of iCCA for the purpose of evaluating the sensitivities of patients in the cluster 1 and cluster 2, or in the low- and high-risk groups to these common chemotherapy drugs. The IC50 values of common chemotherapy drugs for each iCCA patient were calculated (**Figure S5)**. Interestingly, we have found that there was several difference in IC50 values of trametinib, X17-ACC, CI-1040, PD-0325901 and gemcitabine between cluster 1 and cluster 2 showed significant differences (∣logFC∣>0.5) (Fig.8A). The IC50 values of these five drugs are significantly different between two different clusters (p<0.001, Fig.8B). Meanwhile, a significant difference in IC50 values of trametinib, CI-1040, X17-ACC, and PD-0325901 could been observed between high- and low-NPS risk score subgroups (∣logFC∣>0.5) (Fig.8C). Similarly, IC50 values of these four drugs were significantly negatively correlated with the NPS-risk score (Fig.8D). In a nutshell, it is evident from these results that NRGs are essential for the sensitivity of drugs in iCCA.

**Figure 8.**
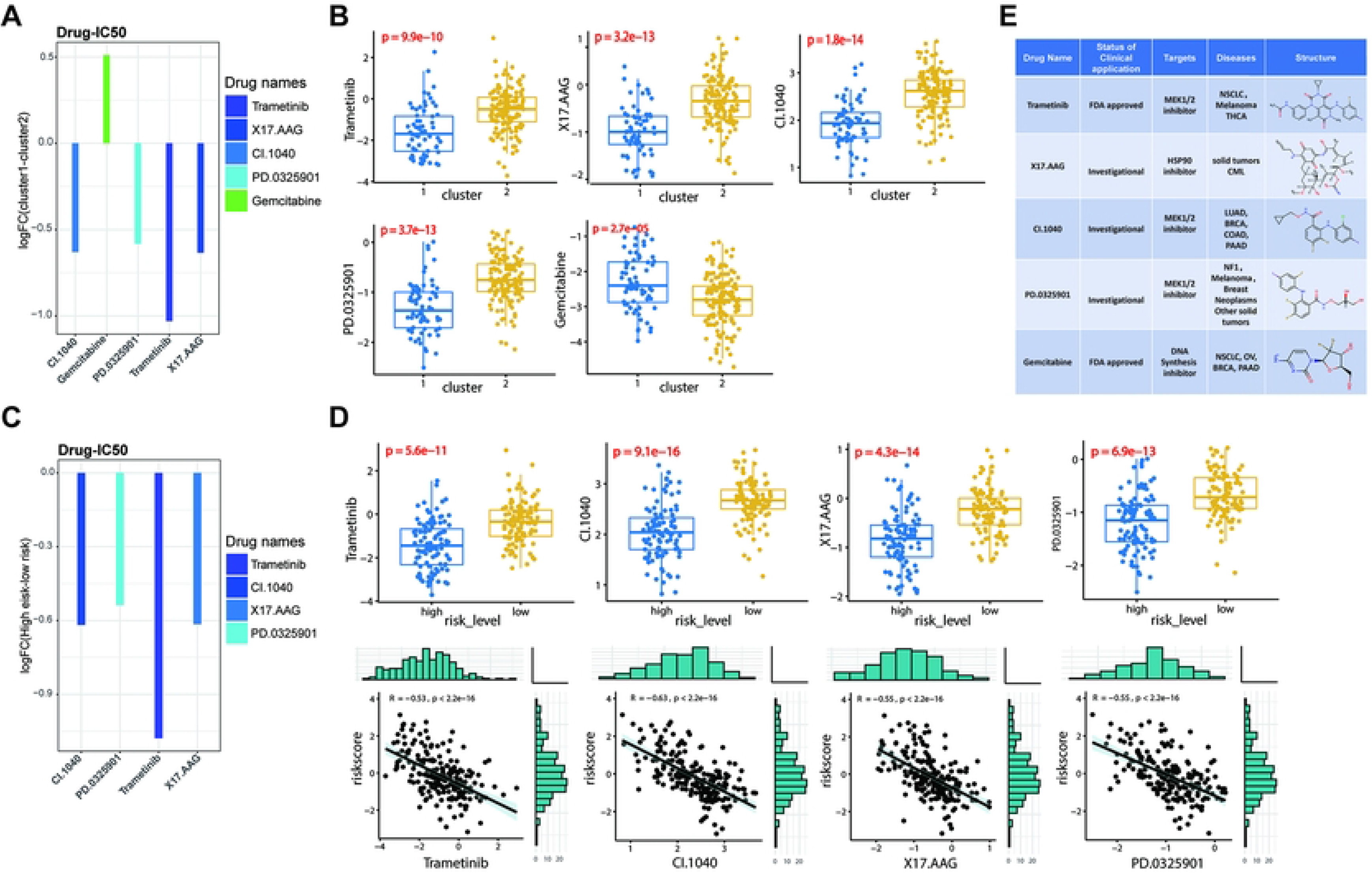
Relationships between NRG-risk score and chemotherapeutic sensitivity. (A and B) The IC50 value of five chemotherapy drugs between cluster 1 and cluster 2 subgroups. ∣logFC∣>0.5. (C-D) The IC50 value of four chemotherapy drugs between high- and low-NPS risk score subgroups. ∣logFC∣>0.5. (E) Summary of status of clinical application, targeted genes, targeted diseases, drug structures of five chemotherapy drugs (trametinib, X17-ACC, CI-1040, PD-0325901 and gemcitabine).

Last but not least, through the investigation of the characteristics and application of these five drugs in iCCA and other tumors, the conclusion that only trametinib and gemcitabine are clinically approved by FDA for the treatment of human cancer were found, while the remaining three drugs are currently in the investigational stage. Moreover, five drugs had not been used in the clinical treatment for iCCA, which also suggests that they can be developed as new drugs for the future treatment of iCCA.

### Analysis of phosphorylation site characteristics of NRGs in iCCA

Due to the results of the previous studies which have demonstrated several significant differences in the enriched pathways, related gene expression between different clusters and high and low risk groups at transcriptional, protein and phosphorylation levels, it is advisable for us to explore the relationship between the different phosphorylation sites of each NRG and patients’ survival. As shown in Fig.9A, all phosphorylation sites of 10 NRG in the TCGA-CHOL cohort were screened, meanwhile, the CHOL patients was divided into high- and low-phosphorylation groups according to the phosphorylation level of every phosphorylation site. Interestingly, there are significant differences in the expression levels of NRGs between the high- and low-phosphorylation groups. Only CHMP28, NF2, CIRBP, PRKACA and CYB5R1 have their own corresponding phosphorylation sites and were all highly expressed in the high phosphorylation level group (all p<0.01; Fig. 9A). By the means of the qPhos database and ActiveDriverDB, the specific positions and sequences of these phosphorylation sites were further retrieved (Fig.9B).

**Figure 9.**
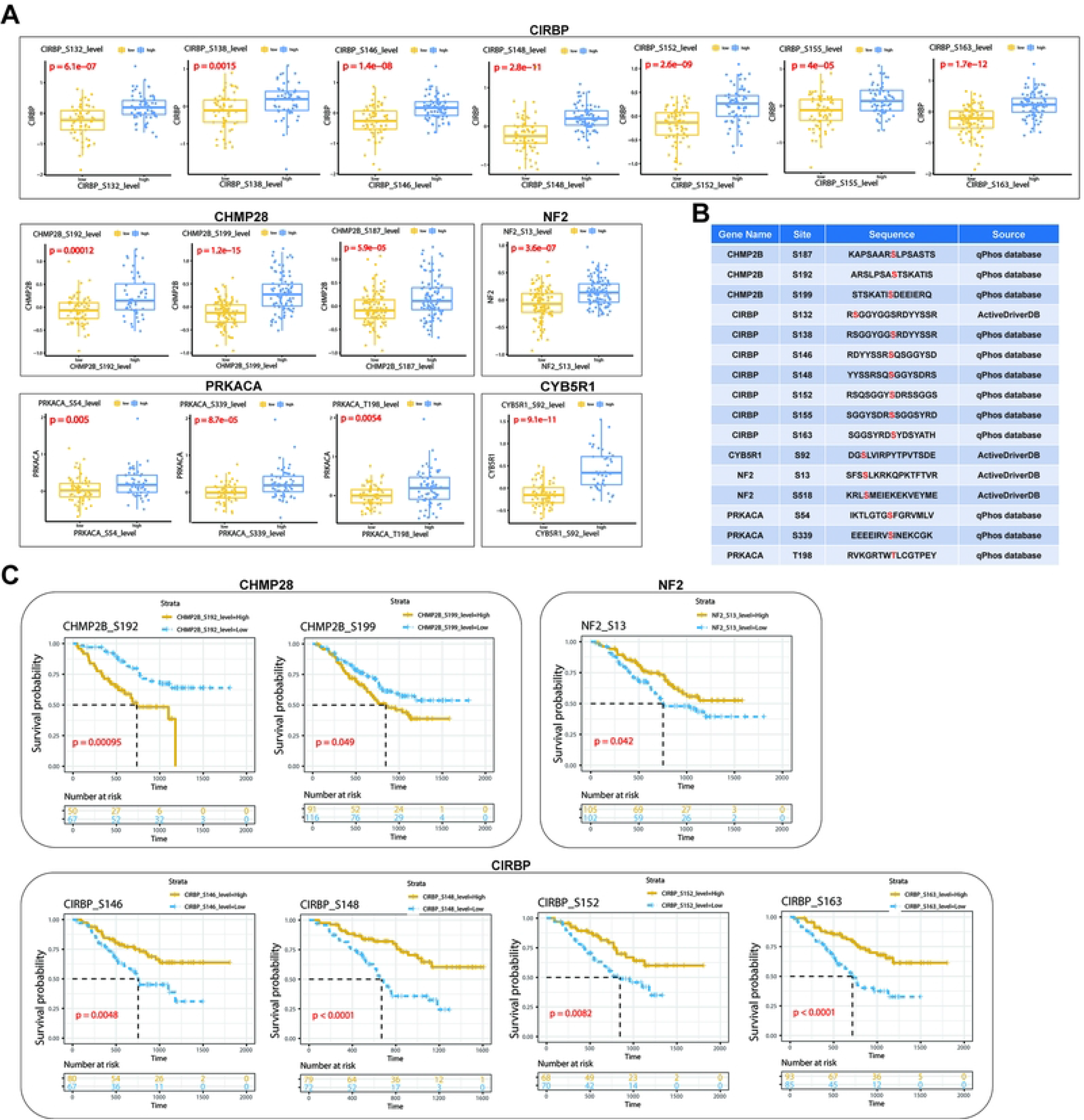
Analysis of phosphorylation site characteristics of NRGs in iCCA. (A) Potential phosphorylated sites levels of CIRBP, CHMP28, NF2, PRKACA and CYB5R1 through the TCGA-CHOL cohort. (B) Potential phosphorylated sites from A and their sequences were predicted by the qPhos database and ActiveDriverDB. All p<0.05. (C) Kaplan-Meier curves showing that the association between phosphorylated sites levels and overall survival in the TCGA-CHOL cohort. All p<0.05.

At the same time, the phosphorylation level of several phosphorylation sites was related to the survival of CHOL patients (Fig.9C). These data indicated that the phosphorylation modification could not only increase the gene expression but also impacted CHOL patients’ survival.

### Expression identification of NRGs in iCCA

Based on TCGA-CHOL cohort, compared with the normal patients (n=9), we found nine of 10 NRGs were up regulated in the CHOL patients (n=36). Only CP was significantly downregulated in CHOL patients (**Fig.10A**). Furthermore, we detected the protein expression of 10 NRGs in 5 pairs of iCCA clinical and adjacent control samples collected from our hospital, and found a similar trend (**Fig.10B**). By the means of the HPA database, it can be found that all 10 NRGs had stronger staining signal in cholangiocarcinoma patient than non-cancer tissues from the representative images of IHC, which is shown in the **Fig.10C**, while there were results suggesting that NRGs may have certain specificity and potential function in regulating tumorigenesis in iCCA.

**Figure 10.** Expression identification of NRGs in iCCA. (A) Heat map showing the expressions of 10 NRGs between tumor and normal patients based on the TCGA-CHOL cohort. (B) The expression of 10 NRGs was observed by Western blot in five pairs of iCCA tissues and adjacent control samples. (C) Representative protein staining images of RAD51 in cholangiocarcinoma cancer and normal tissues. The images were downloaded from the Human Protein Atlas (HPA).

## Discussion

Numerous studies have revealed the indispensable role of nonapoptotic regulatory cell death (NRCD) in innate immunity and antitumor effects ^13,16^. However, most studies have focused on a single NRCD form, which lead that the overall effect and TME infiltration characteristics mediated by the combined effects of multiple NRCD- related genes have not yet been fully elucidated. By the means of the present study, the global alterations in NRGs at the transcriptional and genetic levels in iCCA were revealed. We identified two distinct molecular clusters based on 52 NRGs. Compared to patients with cluster 2, patients in cluster 1 had more advanced clinicopathological features and worse OS. The characteristics of the TME also differed significantly between the two clusters. The NRG-related clusters were also featured by a significant metabolic-activated and immune activation pathways, such as the glycan biosynthesis, mTOR signaling pathway, glycerophospholipid metabolism, MAPK signaling pathway, B cell receptor signaling pathway, nature killer cell mediated cytotoxicity. Furthermore, differences in mRNA transcriptomes between distinct clusters were significantly associated to NRGs and immune-related biological pathways.

Therefore, we constructed the robust and effective NRG risk prognostic signature (NPS) model and demonstrated its predictive ability. Thus, the robust and effective NRG risk prognostic signature (NPS) model was constructed, meanwhile, its predictive ability was demonstrated. The expression levels of 10 NRGs genes included the NPS model in iCCA tissues were also explored. The NRG-related clusters characterized showed lower and higher risk score respectively. Significantly different clinicopathological characteristics, prognosis, mutation, TME, and drug susceptibility were shown by the patients with low- and high-risk scores as well. Finally, by integrating the risk score and 12 clinicopathological characteristics, the quantative nomogram that further improved the performance and facilitated the use of the NRG risk score was established. The prognostic model can be used for prognosis stratification of patients with iCCA, which not only has the ability to assist in better understanding the molecular mechanism of iCCA but also can provide new ideas for targeted therapies.

The development and progression of cancer in concert with alterations in the surrounding stroma. Cancer cells can functionally sculpt their microenvironment through the secretion of various cytokines, chemokines and other factors. This results in a reprogramming of the surrounding cells, enabling them to play a determinative role in tumor survival and progression. The diversity of the tumour microenvironment (TME) of intrahepatic cholangiocarcinoma (iCCA) has not been comprehensively assessed. Although there have been certain advances in immunotherapy recently, patients with iCCA still show heterogeneity in their outcomes, highlighting the crucial role of TME in iCCA tumorigenesis and progression ^24^. Immune cells participate in various immune responses and activities, such as the inflammatory response coordinated by tumors to promote survival ^25^. Evidence has also shown the significant effects of the TME on tumor development, progression, and therapeutic resistance ^26^. Immune cells are important constituents of the tumor stroma and critically take part in this process. Growing evidence suggests that the innate immune cells (macrophages, neutrophils, dendritic cells, innate lymphoid cells, myeloid-derived suppressor cells, and natural killer cells) as well as adaptive immune cells (T cells and B cells) contribute to tumor progression when present in the tumor microenvironment (TME) ^27^ . Cross-talk between cancer cells and the proximal immune cells ultimately results in an environment that fosters tumor growth and metastasis. We discovered that the characteristics of the TME and the relative abundance of 24 immune cells differed significantly between the two NRG-related clusters and different NRG risk-score groups. This finding has suggested the critical role of NRGs in iCCA progression. Increasing evidence has shown that effector CD4+ T cells, playing a vital role in the immune defense of iCCA ^28-30^. Tumor-infiltrating CD4 regulatory T cells exhibited highly immunosuppressive characteristics ^28^. Cluster 2 and low NRG risk score which had a better prognosis, showed higher infiltration of activated CD4+ T cells, demonstrating that CD4+ T cells play a positive role in iCCA development. In the low- NPS- risk socre subgroup, there have been a frequent communication between CD4+ T cells and the other five immune cells which is relatively high, but the communication intensity of common communication pairs between CD4 +T cells and other immune cells was significantly increased in the high-NPS-risk score subgroup, such as MIF-CD74+CXCR4, MIF-CD74+CD44, MDK-SDC1/2/4+NCL and VEGFA/B-VEGFR1. In the TME of iCCA, the complex communication relationship of immune cells was reflected via these data. Although the evidence we got has shown that CD4+ T cells may be the key cells of immune cell communication, its relationship with the NPS-risk score is ambiguous and needs further experimental verification.

We further selected the erythroid cells that communicate most closely with CD4+T cells to explore the molecular mechanism of communication between them. Our study has integrated scRNA-seq of iCCA to identify that *CP* as one of the signatures of high expression in CD4+T cells, suggesting *CP* was potentially involved in tumor evolution.

According to the clustering characteristics of TGF-β expression in erythroid cells, we divided erythroid cells into TGF-β+ and TGF-β- subtypes. And the results indeed showed that CD4+T cells are rich in *CP*, which is also a secreted protein ^31^. The mode of communication between CD4+T cells and TGF-β+ erythroid cells is most likely achieved through the exchange of *CP* and the cytokine TGF-β. …Our findings coincide with the results of a recent study that CD71^+^ erythroid cells (CECs) produce reactive oxygen species to decrease T-cell proliferation. CECs also secrete cytokines, the TGF-β which has the ability to promote T-cell differentiation into regulatory T-cells inclued ^32^. Generally, immune activited-erythroid cells, which possessed dual erythroid and immune regulatory networks, showed immunomodulatory functions while interacted more frequently with various innate and adaptive immune cells ^33^. A study identified erythropoietin and erythroid cells as new participants in tumor-host interactions, while highlighted the involvement of multiorgan signaling events in their induction in response to environmental stress and tumor growth ^34^. These evidences have suggested that the molecular mechanism of communication between CD4+T cells and erythroid cells can be used as a new entry point for the study of iCCA immunotherapy.

The inhibition of immunoinhibitory molecules such as PD-1 and PD-L1 can lead to tumor regression by restoring the cytotoxicity of immune cells ^35^. Pembrolizumab, a PD-L1 inhibitor, has been approved by the Food and Drug Administration (FDA) for the treatment of patients with metastatic or unresectable tumor DNA mismatch repair (MMR) deficiency and/or microsatellite instability (MSI) -high solid tumors after initial therapy, which would include those with cholangiocarcinoma ^36^. Nivolumab (Opdivo), a kind of PD-1-binding IgG4 immunoglobulin, which has shown activity against a wide spectrum of advanced cancers, acts as an immune checkpoint inhibitor by means of blocking the interaction between PD-1 expressed on activated T cells selectively ^37^.

The responses of patients to immune checkpoint inhibitors (ICIs) therapy vary greatly, with some iCCA patients experiencing complete remission while others showing continuous progression ^38^. Here, we showed that there was a significant association between the NRG risk score and the response of iCCA to ICI therapy, while the high-risk score implied the increased sensitivity to ICI, neoadjuvant and adjuvant chemotherapy, which demonstrated that the application of the NRG risk score could assist in decision making for the treatment of iCCA.

In brief, our analysis indicates that the NPS model is an independent risk factor for iCCA, thereby it is obliged to provide an ideal predictor for the prognosis and therapeutic response of iCCA patients. One of the limitations of our study is that the stability of the NRG risk score was tested and validated in a limited number of two independent cohorts and only one scRNA-seq dataset. All analyses were based on data from public databases only, and all samples used in our study were obtained retrospectively. Therefore, an inherent case selection bias may have influenced the results. To prove the reliability of the NRG-related gene signature, studies involving prospective cohorts are needed. In addition, scRNA-seq, a state-of-the-art technology, should be further integrated for future analysis to address possible differences in tumor heterogeneity, immune cell infiltration and intercellular communication between cluster 1 and cluster 2 at single-cell resolution. Moreover, both in vitro and in vivo experiments should be conducted on the discovered NRGs for an in-depth characterization of the mechanisms underlying NRCD regulation and the progression of iCCA in the future.

## Declarations

### Ethics approval and consent to participate

The study was permitted by the Ethics Committee of Shandong Provincial Hospital Affiliated to Shandong University.

### Consent for publication

All authors who have contributed to the study agree to publish it.

### Data Availability

The datasets presented in this study can be found and downloaded from GEO (http://www.ncbi.nlm.nih.gov/geo/).

### Code availability

Not applicable.

### Authors’ contributions

Weibing Wang take responsibility for all aspects of the reliability and freedom from bias of the data presented and their discussed interpretation, drafting the article. Shifeng Xu take responsibility for complete text evaluation and guidance, final approval of the version to be submitted. All authors read and approved the final manuscript.

## Acknowledgments

Not applicable.

## Supplementary Figures

**Figure S1. Creating and evaluating a nomogram.**

**(A)** The nomogram for predicting the 1/2/3-year OS of iCCA patients based on adjuvant therapies, TNM stage, glutamyl transferase, ALT aminoleucine transferase, CEA carcinoembryonic antigen, CA19 carbohydrate antigen, preoperative serum AFP, ALB albumin, tumor size diameter, HBsAg, age, sex and risk score.

**(B)** Calibration plots of the nomogram for predicting the probability of OS at 1-, 2-, and 3-year in the FU-iCCA cohort. The Y-axis represents actual survival, and the X-axis represents nomogram-predicted survival.

**(C-D)** Forest plot of multivariate cox regression analysis for 10 core prognostic NRGs in the (C) GSE78220 melanoma cohort and (D) IMvigor210 cohort, respectively.

**Figure S2. (A-B)** Bar graph showed the proportion of patients with different subtypes of 12 clinical characteristics in (A) cluster 1 and cluster 2 high or in (B) low-NPS risk score subgroups based on FU-iCCA cohort.

**Figure S3. Biological properties and immune landscape of different subgroups based on the FU-iCCA cohort.**

**(A-B)** GSVA analyzed the biological pathways (GO terms and KEGG pathway) of two distinct clusters.

**(C-D)** GSVA analyzed the biological pathways (GO terms and KEGG pathway) of high- and low- NPS risk score subgroups.

**(E)** The landscape of 38 immune cell infiltration between two cluster 1 and cluster 2.

**(F)** The landscape of 38 immune cell infiltration between high- and low- NPS risk score subgroups.

**(G)** StromalScore, ImmuneScore and ESTIMATEScore between two cluster 1 and cluster 2.

**(H)** StromalScore, ImmuneScore and ESTIMATEScore between high- and low- NPS risk score subgroups.

**Figure S4.** t-SNE plots of 10 NRGs expressions in 6 main cell subtypes.

**Figure S5.** Kaplan-Meier curves showing that the association between phosphorylated sites levels and overall survival in the TCGA-CHOL cohort. All p>0.05.

**Figure S6. (A-B)** The IC50 value of all chemotherapy drugs between (A) cluster 1 and cluster 2 subgroups or (B) between high- and low-NPS risk score subgroups.

## References

1. El-Diwany R, Pawlik TM, Ejaz A. Intrahepatic Cholangiocarcinoma. Surgical oncology clinics of North America. Oct 2019;28(4):587–599. doi:10.1016/j.soc.2019.06.002

2. Endo I, Gonen M, Yopp AC, et al. Intrahepatic cholangiocarcinoma: rising frequency, improved survival, and determinants of outcome after resection. Annals of surgery. Jul 2008;248(1):84–96. doi:10.1097/SLA.0b013e318176c4d3

3. Saleh M, Virarkar M, Bura V, et al. Intrahepatic cholangiocarcinoma: pathogenesis, current staging, and radiological findings. Abdominal radiology (New York). Nov 2020;45(11):3662–3680. doi:10.1007/s00261-020-02559-7

4. Han C, Wu T. Cyclooxygenase-2-derived prostaglandin E2 promotes human cholangiocarcinoma cell growth and invasion through EP1 receptor-mediated activation of the epidermal growth factor receptor and Akt. The Journal of biological chemistry. Jul 17 2015;290(29):17806. doi:10.1074/jbc.A115.500562

5. Wink DA, Grisham MB, Mitchell JB, Ford PC. Direct and indirect effects of nitric oxide in chemical reactions relevant to biology. Methods in enzymology. 1996;268:12–31. doi:10.1016/s0076-6879(96)68006-9

6. Churi CR, Shroff R, Wang Y, et al. Mutation profiling in cholangiocarcinoma: prognostic and therapeutic implications. PloS one. 2014;9(12):e115383. doi:10.1371/journal.pone.0115383

7. Zou S, Li J, Zhou H, et al. Mutational landscape of intrahepatic cholangiocarcinoma. Nature communications. Dec 15 2014;5:5696. doi:10.1038/ncomms6696

8. Duan WJ, He RR. Cuproptosis: copper-induced regulated cell death. Sci China Life Sci. Aug 2022;65(8):1680–1682. doi:10.1007/s11427-022-2106-6

9. Galluzzi L, Vitale I, Aaronson SA, et al. Molecular mechanisms of cell death: recommendations of th e Nomenclature Committee on Cell Death 2018. Cell Death Differ. Mar 2018;25(3):486–541. doi:10.1038/s41418-017-0012-4

10. Carneiro BA, El-Deiry WS. Targeting apoptosis in cancer therapy. Nat Rev Clin Oncol. Jul 2020;17(7):395–417. doi:10.1038/s41571-020-0341-y

11. Mishchenko TA, Balalaeva IV, Vedunova MV, Krysko DV. Ferroptosis and Photodynamic Therapy Synergism: Enhancing Anticancer Treatment. Trends Cancer. Jun 2021;7(6):484–487. doi:10.1016/j.trecan.2021.01.013

12. Gu Z, Liu T, Liu C, et al. Ferroptosis-Strengthened Metabolic and Inflammatory Regulation of Tumor-Associated Macrophages Provokes Potent Tumoricidal Activities. Nano Lett. Aug 11 2021;21(15):6471–6479. doi:10.1021/acs.nanolett.1c01401

13. Zeng C, Lin J, Zhang K, et al. SHARPIN promotes cell proliferation of cholangiocarcinoma and inhibits ferroptosis via p53/SLC7A11/GPX4 signaling. Cancer science. Nov 2022;113(11):3766–3775. doi:10.1111/cas.15531

14. Qiao Y, Choi JE, Tien JC, et al. Autophagy Inhibition by Targeting PIKfyve Potentiates Response to Immune Checkpoint Blockade in Prostate Cancer. Nat Cancer. Sep 2021;2:978–993. doi:10.1038/s43018-021-00237-1

15. Xie Y, Zhu S, Zhong M, et al. Inhibition of Aurora Kinase A Induces Necroptosis in Pancreatic Carcinoma. Gastroenterology. Nov 2017;153(5):1429–1443 e5. doi:10.1053/j.gastro.2017.07.036

16. Song W, Ren J, Xiang R, Kong C, Fu T. Identification of pyroptosis-related subtypes, the development of a prognosis model, and characterization of tumor microenvironment infiltration in colorectal cancer. Oncoimmunology. 2021;10(1):1987636. doi:10.1080/2162402x.2021.1987636

17. Dong L, Lu D, Chen R, et al. Proteogenomic characterization identifies clinically relevant subgroups of intrahepatic cholangiocarcinoma. Cancer cell. Jan 10 2022;40(1):70–87.e15. doi:10.1016/j.ccell.2021.12.006

18. Mariathasan S, Turley SJ, Nickles D, et al. TGFβ attenuates tumour response to PD-L1 blockade by contributing to exclusion of T cells. Nature. 2018/02/01 2018;554(7693):544-548. doi:10.1038/nature25501

19. Tsvetkov P, Coy S, Petrova B, et al. Copper induces cell death by targeting lipoylated TCA cycle proteins. Science. Mar 18 2022;375(6586):1254-1261. doi:10.1126/science.abf0529

20. Zhou N, Bao J. FerrDb: a manually curated resource for regulators and markers of ferroptosis and ferroptosis-disease associations. Database : the journal of biological databases and curation. Jan 1 2020;2020doi:10.1093/database/baaa021

21. Colaprico A, Silva TC, Olsen C, et al. TCGAbiolinks: an R/Bioconductor package for integrative analysis of TCGA data. Nucleic acids research. 2016;44(8):e71–e71. doi:10.1093/nar/gkv1507

22. Mayakonda A, Lin DC, Assenov Y, Plass C, Koeffler HP. Maftools: efficient and comprehensive analysis of somatic variants in cancer. Genome research. Nov 2018;28(11):1747–1756. doi:10.1101/gr.239244.118

23. Wilkerson MD, Hayes DN. ConsensusClusterPlus: a class discovery tool with confidence assessments and item tracking. Bioinformatics. 2010;26(12):1572–1573. doi:10.1093/bioinformatics/btq170

24. Martin-Serrano MA, Kepecs B, Torres-Martin M, et al. Novel microenvironment-based classification of intrahepatic cholangiocarcinoma with therapeutic implications. Gut. May 18 2022;doi:10.1136/gutjnl-2021-326514

25. Seager RJ, Hajal C, Spill F, Kamm RD, Zaman MH. Dynamic interplay between tumour, stroma and immune system can drive or prevent tumour progression. Convergent Science Physical Oncology. 2017/07/28 2017;3(3):034002. doi:10.1088/2057-1739/aa7e86

26. Hinshaw DC, Shevde LA. The Tumor Microenvironment Innately Modulates Cancer Progression. Cancer research. 2019;79(18):4557–4566. doi:10.1158/0008-5472.CAN-18-3962

27. Turley SJ, Cremasco V, Astarita JL. Immunological hallmarks of stromal cells in the tumour microenvironment. Nature Reviews Immunology. 2015/11/01 2015;15(11):669-682. doi:10.1038/nri3902

28. Zhang M, Yang H, Wan L, et al. Single-cell transcriptomic architecture and intercellular crosstalk of human intrahepatic cholangiocarcinoma. Journal of hepatology. Nov 2020;73(5):1118–1130. doi:10.1016/j.jhep.2020.05.039

29. Tran E, Turcotte S, Gros A, et al. Cancer immunotherapy based on mutation-specific CD4+ T cells in a patient with epithelial cancer. Science. May 9 2014;344(6184):641-5. doi:10.1126/science.1251102

30. Song G, Shi Y, Meng L, et al. Single-cell transcriptomic analysis suggests two molecularly subtypes of intrahepatic cholangiocarcinoma. Nature communications. Mar 28 2022;13(1):1642. doi:10.1038/s41467-022-29164-0

31. de Oliveira AL, Gallo M, Pazzagli L, et al. The structure of the elicitor Cerato-platanin (CP), the first member of the CP fungal protein family, reveals a double ψβ-barrel fold and carbohydrate binding. The Journal of biological chemistry. May 20 2011;286(20):17560–8. doi:10.1074/jbc.M111.223644

32. Grzywa TM, Nowis D, Golab J. The role of CD71(+) erythroid cells in the regulation of the immune response. Pharmacology & therapeutics. Dec 2021;228:107927. doi:10.1016/j.pharmthera.2021.107927

33. Xu C, He J, Wang H, et al. Single-cell transcriptomic analysis identifies an immune-prone population in erythroid precursors during human ontogenesis. Nature immunology. Jul 2022;23(7):1109–1120. doi:10.1038/s41590-022-01245-8

34. Sano Y, Yoshida T, Choo MK, et al. Multiorgan Signaling Mobilizes Tumor-Associated Erythroid Cells Expressing Immune Checkpoint Molecules. Molecular cancer research : MCR. Mar 2021;19(3):507–515. doi:10.1158/1541-7786.Mcr-20-0746

35. Bellmunt J, Powles T, Vogelzang NJ. A review on the evolution of PD-1/PD-L1 immunotherapy for bladder cancer: The future is now. Cancer treatment reviews. 2017;54:58–67. doi:10.1016/j.ctrv.2017.01.007

36. Naganuma A, Sakuda T, Murakami T, et al. Microsatellite Instability-high Intrahepatic Cholangiocarcinoma with Portal Vein Tumor Thrombosis Successfully Treated with Pembrolizumab. *Internal medicine (Tokyo*, Japan). Sep 15 2020;59(18):2261–2267. doi:10.2169/internalmedicine.4588-20

37. Fontugne J, Augustin J, Pujals A, et al. PD-L1 expression in perihilar and intrahepatic cholangiocarcinoma. Oncotarget. Apr 11 2017;8(15):24644–24651. doi:10.18632/oncotarget.15602

38. Rizzo A, Ricci AD, Brandi G. Recent advances of immunotherapy for biliary tract cancer. Expert review of gastroenterology & hepatology. May 2021;15(5):527–536. doi:10.1080/17474124.2021.1853527

